# Taking stock of selective logging in the Andaman Islands, India: recent & legacy effects of timber extraction, assisted natural regeneration and a revamped working plan

**DOI:** 10.1101/2024.06.23.600264

**Authors:** Akshay Surendra, Vanjulavalli Sridhar, Anand M. Osuri, Jayashree Ratnam

## Abstract

Forest management is an evolving balance between biodiversity conservation and economic needs. A shift in Andaman Islands’ Working Plan mandate in 2000s reflects this evolution. Our study independently assesses the impact of said policy change on post-logging recovery of forests in Baratang and Middle Andaman.

In 2017-18, we placed seventy-six 0.49ha plots across evergreen and deciduous patches and compared large-tree (≥180cm girth) density and diversity across forests that were logged after 2005 focussing on sustainability, logged in 1990s focussing on timber, logged twice in 1990s and after 2005, and unlogged forests. We assessed forest regeneration in thirty 0.01ha plots along a coupe road within forests logged after 2005.

Forests logged after 2005 had similar density of large trees as forests logged in 1990s (despite having 1/3^rd^ the recovery time), indicating reduced offtake or better recruitment. Along the unlogged—once-logged—twice-logged gradient, the dominance of *Pterocarpus dalbergioides* in deciduous patches decreased while the dominance of *Diptercarpus sp.* in evergreen patches increased. Compared to natural regeneration, proportionately more deciduous saplings were planted in both evergreen and deciduous patches.

The new working plan maintains timber stock but not diversity. We make six simple recommendations to better align practice with the Working Plan mandate.

**Synthesis:** Post-2005 timber extraction policy in the Andaman Islands is partially successful but long-term forest health, in line with the working plan mandate, requires (1) lower timber offtake from deciduous patches and (2) targeted assisted regeneration of non-timber tree species.

## INTRODUCTION

Selective logging is a pervasive, widespread anthropogenic disturbance in forests around the world (Malhi et al. 2014). The widespread scale of selective logging is at least in part because it is seen as the least destructive of disturbances (Gibson et al. 2011), often promoted as a possible middle-ground that balances biodiversity conservation and economic demand (Chaudhary et al. 2016). However, there has been no systematic test of whether past and current logging regimes achieve this middle ground.

In the Andaman Islands of India and elsewhere in the tropics, an ideal middle-ground hinges on two critical parameters that determine the impact of selective logging on forest structure and recovery: intensity of logging and frequency of logging. The intensity of selective logging – indicated by the number of trees harvested from a patch of forest and the extent of damage to smaller trees during logging operations – needs to be high enough for economic viability but low enough so that timber stocks can recover. Similarly, the frequency of logging must by low enough that there is enough time for forests to recover, but frequent enough to assure sustained monetary benefit. Practices like reduced-impact logging or caps on maximum extraction (< 8 trees/ha) modulate intensity of logging (Putz et al. 2008). However, the search for an optimal logging frequency is more elusive - empirical and modelling studies tell us that selective logging beyond three cycles becomes unviable (Zimmerman and Kormos 2012).

A long history of logging over 150-200 years in the Andaman Islands (Bhattee 1958) has allowed ecological constraints to catch up with financial sustainability of timber harvest. Large-scale selective logging took off with the invention of the Andaman Shelterwood System (Chengappa 1934): between the 1950s and 1990s, the Andaman Islands Forest Department extracted 100,000 – 150,000 cubic metres of timber annually (Chaudhry 1998), with private logging companies in Diglipur and Mayabunder regions extracting 2-3 times as much timber. By the 1990s, two out of three private logging companies had shut down their sawmills, citing insufficient timber (Sekhsaria 2004 and references therein). The Forest Department too had reduced its extraction quota on its own volition by the late 1990s, recognizing its role as “guardians of natural riches of the Islands and not as a source of revenue” (Chaudhry 1998). Moreover, the Jarawa Tribal Reserve on South Andaman, Baratang and Middle Andaman Islands had been periodically reduced in size to access unlogged forests within the tribal reserve for timber, hinting at a reduction in timber stocks elsewhere in the archipelago (forests that were “returned” to tribal reserve status after timber was harvested - Sekhsaria and Pandya 2010 and references therein). Alarmingly, the Forest Department even began to harvest timber from within the Onge Tribal Reserve on the island of Little Andaman (Sekhsaria 2004), significantly further south and isolated from the forest operations on the central group of islands (North, Middle and South Andaman). All these lines of evidence point to a scenario of severely depleted timber stocks across the archipelago by the 1990s.

But what caused such a depletion of timber stocks? Modelling studies concede that the highest revenue from timber is always from unlogged forests: 40 years after a logging event, timber value is still only ∼38% of what an unlogged forest would contain (Fisher et al. 2011; Putz et al. 2012). Given the large volumes of timber extracted and the high demand from mainland India (98% of extracted timber from the Andaman Islands in the 1990s were used in mainland India: Sekhsaria 2004), it is very likely that a large proportion of timber extracted from the Andaman Islands came from previously unlogged forests. Older members of the local forest department staff concurred with this assessment, citing a preference for “*khara*” (unlogged) jungle. When unlogged forests became scarce, and timber from logged forests wasn’t recovering quickly enough, timber stocks declined. By 2002, when the Supreme Court of India stepped in to revamp timber operations in the Andaman Islands, the Forest Department had already responded to the dwindling timber supply by reducing its volume targets by half to ∼40,000 m^3^(Sekhsaria 2004).

The Supreme Court intervened in response to a public interest litigation (first filed in the Calcutta High Court), temporarily stopped all logging activity, set up the Shekhar Singh committee to look into the matter and later accepted its recommendations (Sekhsaria 2004). The Committee and the judiciary took a holistic view, identifying felling in the Onge Tribal Reserve as symptomatic of a deeper problem of unsustainable timber extraction. Broadly, the Shekhar Singh Committee recommended a shift in policy from maintaining sustainable flows and stocks of timber to conserving biodiversity and livelihoods.

This paradigm shift was reflected in the new working plan starting in 2005 (Singh 2003). The new working plan restricts all future logging to previously logged forests, implicitly conceding that unlogged forests are inviolate. Within previously logged forests, logging intensity was capped at 3 trees/ha per logging event. Logging frequency was fixed at 90 years, where every 30 years, 1/3rd of the available timber would be extracted within the cap of 3 trees/ha or less (Singh 2003). This mandate is a clear departure from higher logging intensities (∼8 trees/ha on average) and shorter logging cycles (35-45 years) in regions where only economic considerations hold sway. On the demand side, the export of raw timber to mainland India was banned; only finished products could now be exported outside the Andaman Islands. To enforce compliance in the long term, hand-drawn coupe maps were replaced by boundaries drawn with a GPS, boundaries that delimited logging cycles for the next 90 years. Importantly, previously logged forests was renamed Eco-Restoration Working Circle and the phasing-out of timber plantations (mainly teak) was renamed Plantation Reclamation Circle, highlighting a change in ethos that shifts focus away from timber extraction and towards forest restoration and reclamation (Singh 2003). Explicit recognition of ecological integrity through eco-restoration has important consequences for management, including but not limited to species selection, site selection, strategies of planting over time, as well as strategies to effectively complement active (assisted regeneration) restoration with passive (natural regeneration) restoration (Krishnan and Osuri 2023; Bartholomew et al. 2024)

But has this new policy changed how forests are managed on-ground? Our study addresses this question by comparing the impact of logging under this new policy (between 2007 and 2014) aimed at biodiversity conservation and livelihood protection against logging under the old policy (in the 1990s) aimed at timber extraction. We used large remnant trees above logging girth (≥180cm) as a response metric, where large remnant trees are an estimate of timber stock: more timber off-take results in fewer large remnant trees, and vice versa. If the density and diversity of large remnant trees in forests logged in the 1990s (old policy) are similar or lower in density and diversity of large remnant trees within forests logged between 2007-14 (new policy) – despite forests logged in the 1990s having had twice or thrice as much time to recover as those logged in 2007-14 – we can infer that the new policy is an improvement. It is important to note that forests logged new policy had been logged in the 1940s but we treat the 70-80-year time window to be long enough to allow near complete recovery of timber stock (Blanc et al. 2009). We also enumerated large remnant trees in forests logged twice (recently under the new policy and earlier under the old policies) to understand cumulative effects of multiple logging events. Finally, we enumerated large remnant trees in unlogged forests that serve as a control representing the full potential of growing stock. All comparisons of forest recovery under different logging regimes were done separately for ecologically distinct evergreen and deciduous forests. We also assessed the efficacy of tree planting efforts in maintaining or eroding differences between evergreen-deciduous patchiness by comparing the proportion of deciduous and evergreen saplings from assisted versus natural regeneration within a stretch of forest (with both evergreen and deciduous patches) logged under the new policy. In line with a working plan mandate of eco-restoration, we provide a list of local names of trees and palms beyond timber species.

## MATERIAL AND METHODS

This study was conducted in the Andaman archipelago within India’s Andaman and Nicobar Islands union territory. The Andaman Islands are a tropical archipelago within the Indo-Burma biodiversity hotspot with high forest cover (>80%), moderate endemism in flora and fauna, and indigenous (Jarawa, Onge and Great Andamanese) and settler (“Ranchi”, Bengali, Karen, among others) human communities who depend on these forests for direct and indirect benefits (Parkinson 1923; Chengappa 1934; Reddy et al. 2016). These forests have been worked for timber for over 150-200 years, but indigenous people have lived and used these forests for at least 20,000 years (Sekhsaria and Pandya 2010). Our study focuses on the central region of this archipelago: Nilambur and Adajig Ranges within the Baratang Forest Division, and Bajalungta/Kadamtala Range within the Middle Andaman Forest Division.

The Bajalungta Range was logged between 2007 and 2014 (new policy). A major portion of Adajig Range was logged in the 1990s (old policy), the Nilambur range was logged both in the 1990s and the 2007-14 years (new and old policy), while a narrow belt of forest on the North-Western side of Adajig, including the adjoining Island of *Hiran Tikri* and Evergreen Island, were unlogged forests (unlogged control). Each of these four treatments straddled distinct evergreen and deciduous patches (Figure 1 in Surendra et al. 2021).

**Figure 1:**
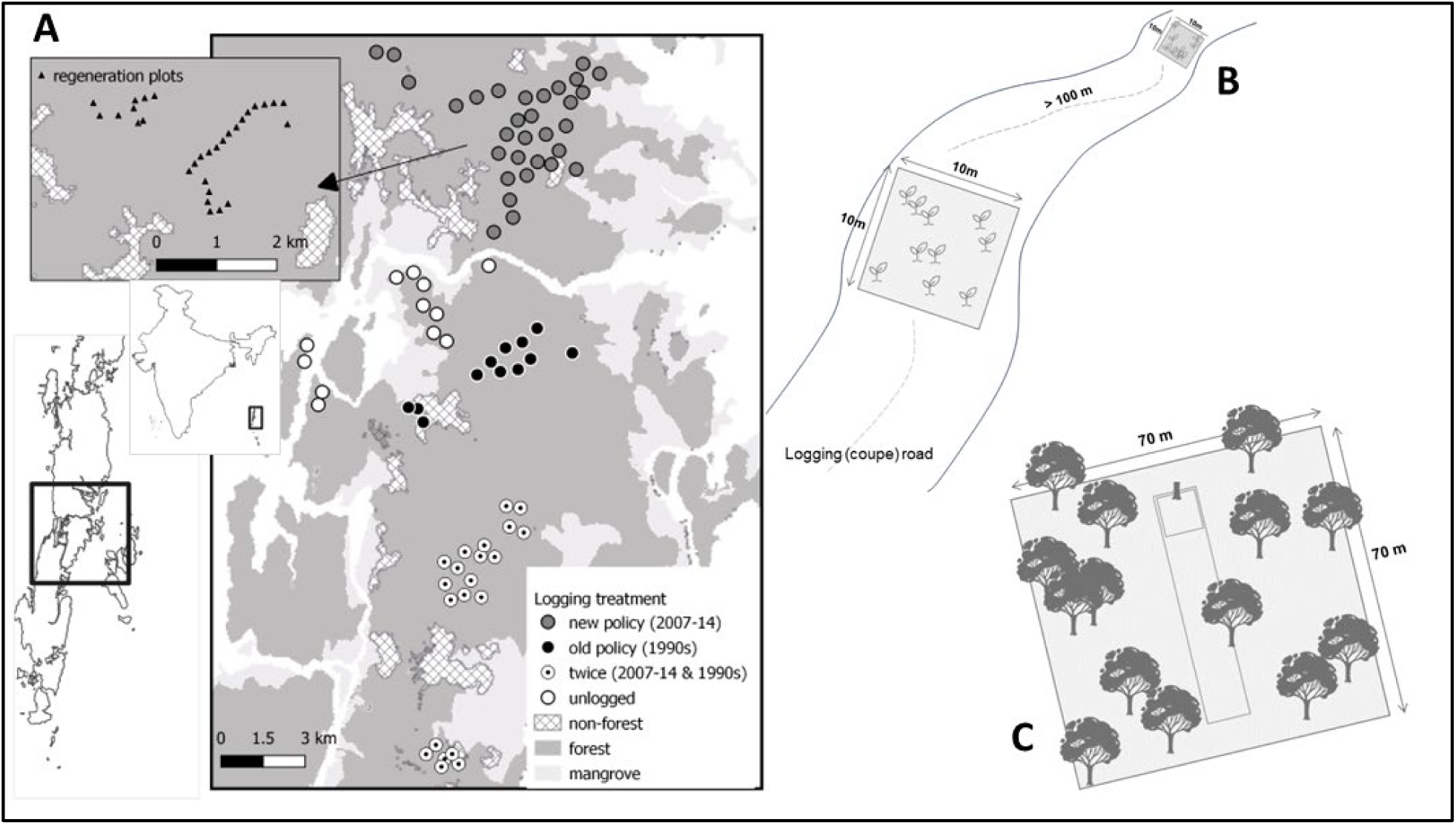
(A) The study site is located within the Middle Andaman and Baratang Forest Divisions of the Andaman and Nicobar Islands of India. The map shows the locations of each of the 76 plots spread across areas logged between 2007-14 (new policy, grey), in the 1990s (old policy, black), logged under both old and new policy (white with a black dot), and unlogged patches (white). The inset plot shows locations of regeneration plots along a coupe road in the Bajalungta range, Middle Andaman. We placed 10x10m plots along the coupe road (B) to enumerate seedlings, and we measured large trees in 70 x 70m plots (C) within which two smaller plots were nested (analysis of nested plot data and the data itself can be found in an earlier study).

To delineate treatments in the field, we obtained new-policy coupe outlines from the current working plans for Baratang and Middle Andaman Forest Divisions and digitized coupe maps under the old policy from photographs of maps in range offices. We avoided uncertainties associated with hand-drawn maps by sampling away from edges of old coupes, with additional guidance from older forest staff members who accompanied us. Such local information was also helpful in identifying unlogged control forests (“*kharajungal*”) and delineating evergreen (“*hara tikri*”) and deciduous patches (“*sukhatikri*”) during the leaf-on wet season.

### Dataset 1: Seventy-six 0.49 ha plots enumerating trees 180cm in circumference or above

Between December 2017 and May 2018, we placed 76 plots across unlogged forests, forests logged under the new policy (2007-14), forests logged under the old policy (1990s), and forests logged under both policies (1990s and 2007-14) each of which overlapped evergreen and deciduous patches – we had at least 6 plots in each logging treatment-forest type combination. Each plot was 70m x 70m (0.49 ha) in size and encircled two smaller plots along a treefall line. Data from the smaller plots have been analysed in a previous study (Surendra et al. 2021b) and the data are available open-access on Data Dryad (Surendra et al. 2021a) Within these plots, we enumerated all remnant large trees above 180cm girth to infer standing stock above logging girth. We identified each tree to the finest taxonomic level possible (morphospecies, if not species) and measured their height or the tallest branches using a rangefinder and girth using measuring tape. Our measurement of remnant timber stock is a generous estimate because, in practice, trees at least 210-230 cm in circumference and only a subset of available species with desirable timber and appropriate bole length and shape are harvested. We did not measure timber offtake directly by enumerating logging stumps because stump decay rates differed by location and species, decay was rapid within evergreen patches compared to deciduous patches while the heartwood of Padauk (*Pterocarpus dalbergioides*) stumps regardless of forest type persisted for at least ∼4 decades (some purportedly from the time of Japanese occupation in the 1940s); such old *Padauk* stumps helped delineate logged areas on-ground.

With this dataset, we first calculated the total number of stems and the total number of unique species within each plot. We then scaled these data from 0.49 ha to 1 ha and plotted them as box plot summaries overlain with raw data points. We identified statistically significant differences in density and species richness between logging treatments using non-parametric Kruskal-Wallis tests suitable for unequal sample sizes. We tested these separately within each forest type. Next, we made rank-abundance curves of large trees for each logging treatment-forest type combination by pooling trees across all plots within each logging-treatment—forest-type combination by species. And because each combination had an unequal number of plots and the lowest number of plots was 6, we scaled the relative abundances in each combination down to 2.94 (0.49 × 6) ha. We labelled dominant trees that contributed the top 50% of stems in each combination.

### Dataset 2: Thirty 0.01 ha plots enumerating saplings 30 cm or above in height but less than 10 cm in circumference

Along a coupe road near the Bajalungta range near *aamtikri*, we placed thirty 10 x 10m plots at least 100m apart, with 16 plots in deciduous patches and 14 in evergreen patches. Within each plot, we counted saplings above 30 cm in height and less than 10cm in GBH and identified them to the finest taxonomic level. These saplings included a mix of naturally growing individuals (natural regeneration) and individuals planted by the forest department (assisted regeneration). Saplings grown in the nursery have a characteristic bulge at the root collar that was visible and identified by trained staff, allowing us to make this distinction between assisted and natural regeneration. Saplings grown in the nursery are never more than a year old and planting in annual cycles. Using these data, we calculated the number of regenerating sapling stems in each 0.01 ha plot, splitting the numbers by habitat (whether the stretch of coupe road was located in an evergreen or deciduous patch), regeneration type (natural regeneration or planted saplings, the latter with a distinctive bulge at the base of the stem), and the phenology of the species as adults (evergreen or deciduous habit); 0.99% of saplings whose specific identity (and consequently, their habit) could not be determined, were discarded. We used a conservative estimate of what saplings were classified as assisted regeneration, combining saplings that we were confident was planted (4.5% of saplings) with saplings that we were not sure about (7.8%). Thus, the fraction of natural regeneration we use may be a conservative estimate of actual natural regeneration. We converted these data into percentages, as a fraction of the number of saplings of each regeneration type (natural or assisted) per plot, to compare the evergreen and deciduous fraction among planted and naturally-growing saplings separately among deciduous and evergreen patches; we plot these data as box plots alongside p-values from unpaired t-tests.

All data exploration, statistical analyses and visualization were done in R programming language using the *base*, *utils*, *stats*, *tidyverse*, *ggpubr*, *ggrepel* and *cowplot* packages (R Core Team 2018). All data alongside code to reproduce these plots are available open-access on GitHub (Surendra 2023). Throughout the study period, we also documented and compiled several local Ranchi (Sadri) and scientific names for tree species of value to local people.

## RESULTS AND DISCUSSION

We sampled 1105 large trees of 80 morphospecies within 76 plots and a total sampling area of 37.24 ha. In regeneration plots along a coupe road, we recorded 3026 individuals of 110 morphospecies within 30 plots, sampling an area of 0.3 ha in total. We compiled local names of c.68 morphospecies used as small timber and Non-Timber Forest Produce (NTFP) by local communities.

We found that differences in the density of large trees between forests logged once, under the new policy versus forests logged under the old policy, were not statistically discernible for both evergreen and deciduous forest patches (p > 0.05, Figure 2). However, within deciduous forests, the species richness of remnant large trees was higher in forests logged once under the old policy compared to forests logged once under the new policy (p = 0.035, Figure 2); corresponding differences in evergreen patches were not statistically discernible. Small sample sizes per treatment precluded statistically significant differences (n = 6 to 18 per forest type per treatment). However, median values in the boxplot indicate that the highest densities and number of species of large remnant trees were within unlogged forests, across evergreen and deciduous patches, and forests logged twice had among the lowest median density and richness of large trees (Figure 2).

**Figure 2:**
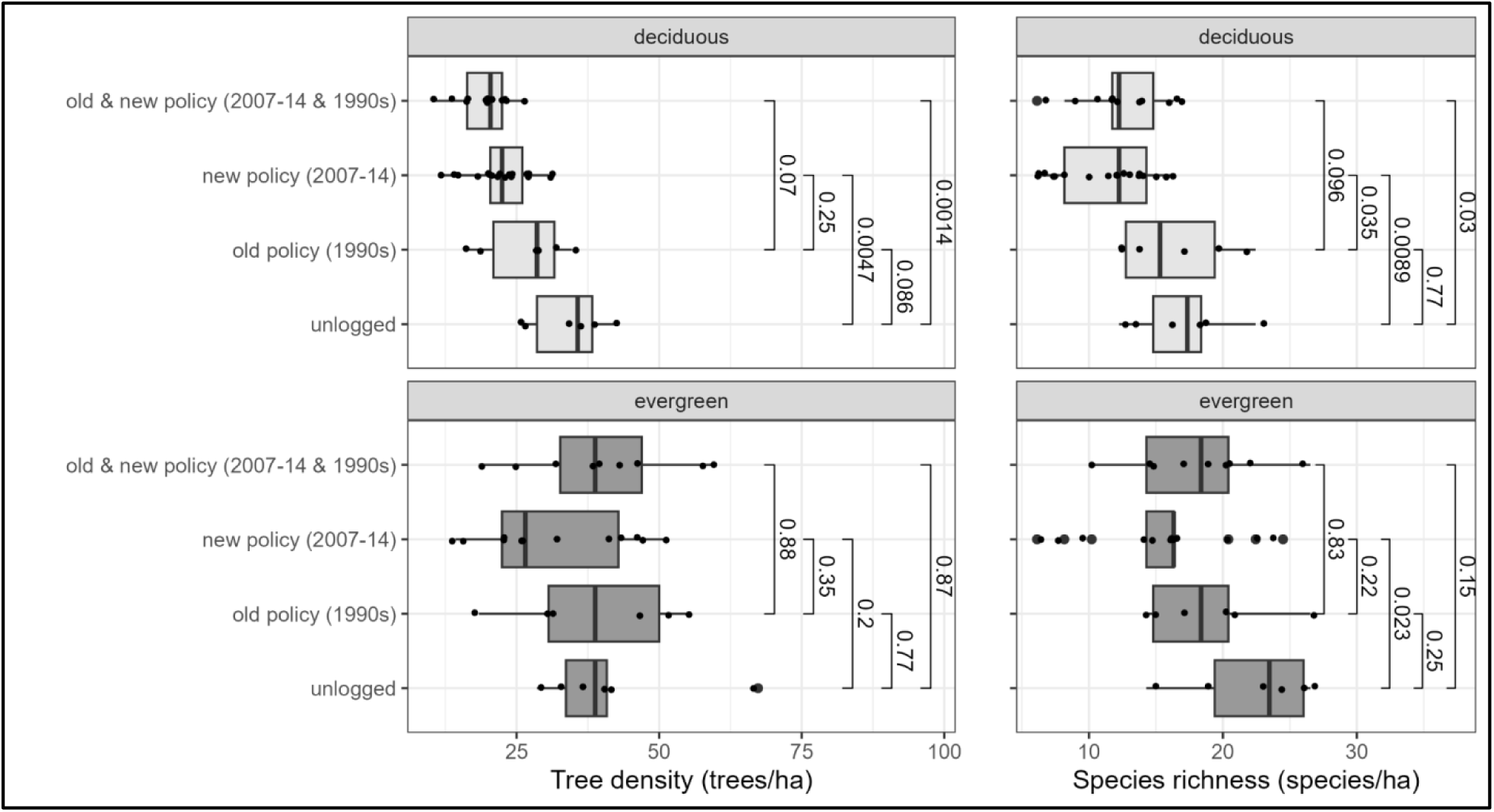
Density (left) & species richness (right) of large trees across variously logged forests and an unlogged control within deciduous (above) and evergreen (below) patches, scaled to 1ha. P-values from statistical tests of differences (non-parametric Kruskal-Wallis test) are shown.

Rank-abundance curves revealed that the most common species were Garjan (*Dipterocarpus sp.*) and Padauk (*Pterocarpus dalbergiodes*) in evergreen and deciduous patches respectively (Figure 3). We find that Padauk density reduced from ∼60 trees/ha in unlogged forests, to ∼40 trees/ha in once-logged forests, to ∼30 trees/ ha in forests logged twice, once under each policy (top row, figure 3). In evergreen patches, the relative dominance of Garjan increases along the unlogged— once-logged—twice-logged gradient, seemingly at the cost of other co-dominants (bottom row, Figure 3).

**Figure 3:**
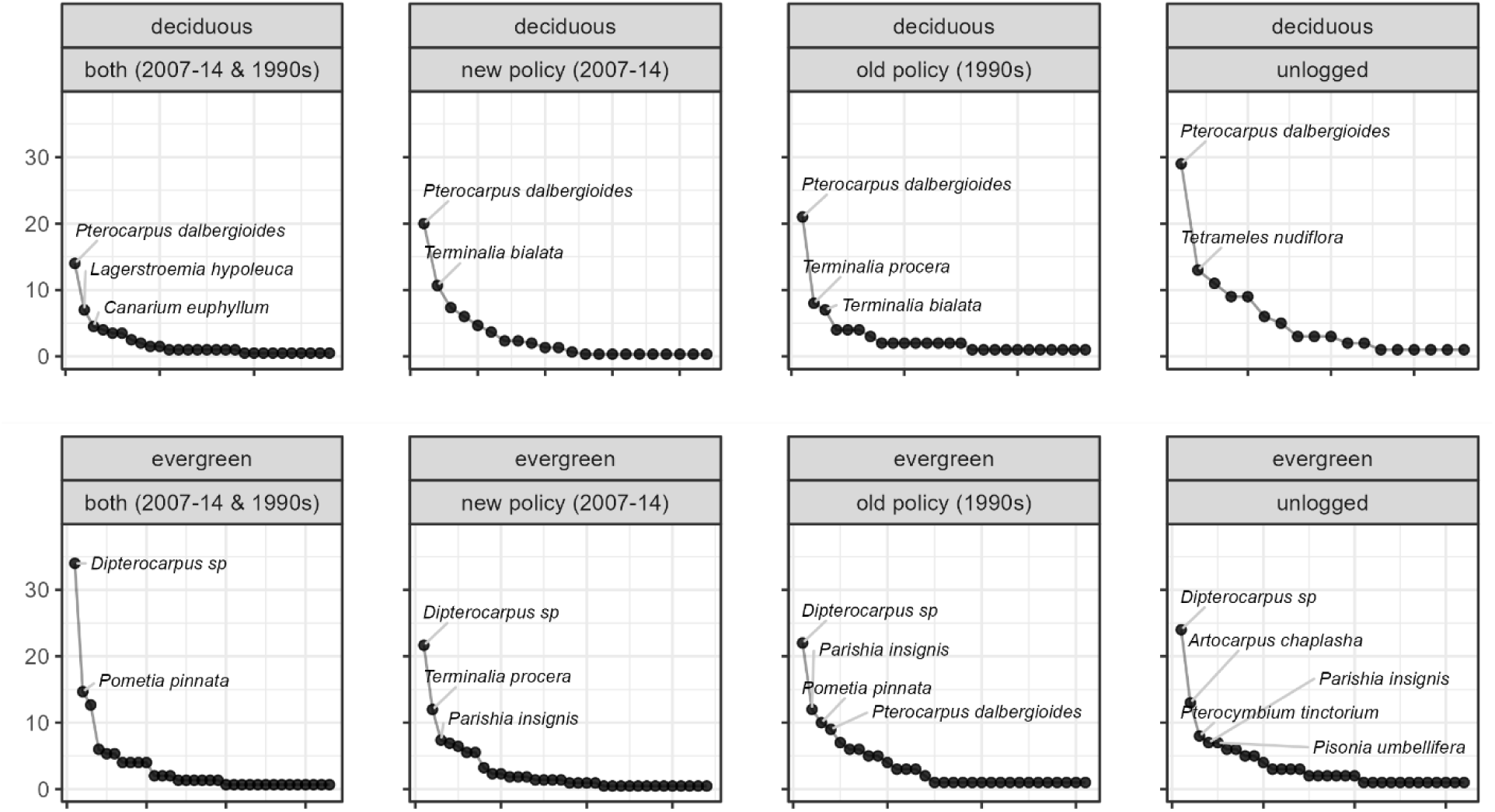
Rank-abundance curves of large trees (>=180 cm DBH) within each logging treatment and forest type. The Y-axis represents abundances scaled to 6 hectares.

Our data on regeneration showed that a greater fraction of seedlings that established naturally were evergreen than deciduous, in both evergreen (p < 0.001) and deciduous patches (p = 0.001) (Figure 4). However, among the saplings that were planted (assisted regeneration), the fraction of saplings of deciduous species that established was higher in deciduous patches (p=0.025). On the other hand, the fractions of saplings of evergreen species that established was not statistically different between evergreen and deciduous patches (p=0.239) (Figure 4). Moreover, the number of saplings growing naturally per unit area was 8.1 times higher than the number of planted saplings. These data on natural regeneration suggest that despite early establishment success, evergreen species regenerating in deciduous patches may be outcompeted. It is likely that deciduous patches cannot support evergreen canopy trees in the dry season without underground water sources (Parkinson 1923; Chengappa 1934). Further, deciduous patches correlate with volcanic soil with Nickel (Pal et al. 2005) that promote dominance of putative Nickel accumulators such as *Rinorea benghalensis*, *Celtis sp.*, *Streblus sp.* and *Dichapetalum gelonoides* (Pal and Paul 2007) that are all restricted to deciduous patches. Together, these environmental filters may filter out evergreen tree species from the canopy of deciduous patches. The assisted regeneration of deciduous saplings and their subsequent increases in establishment in deciduous patches, may further exacerbate this pattern.

**Figure 4:**
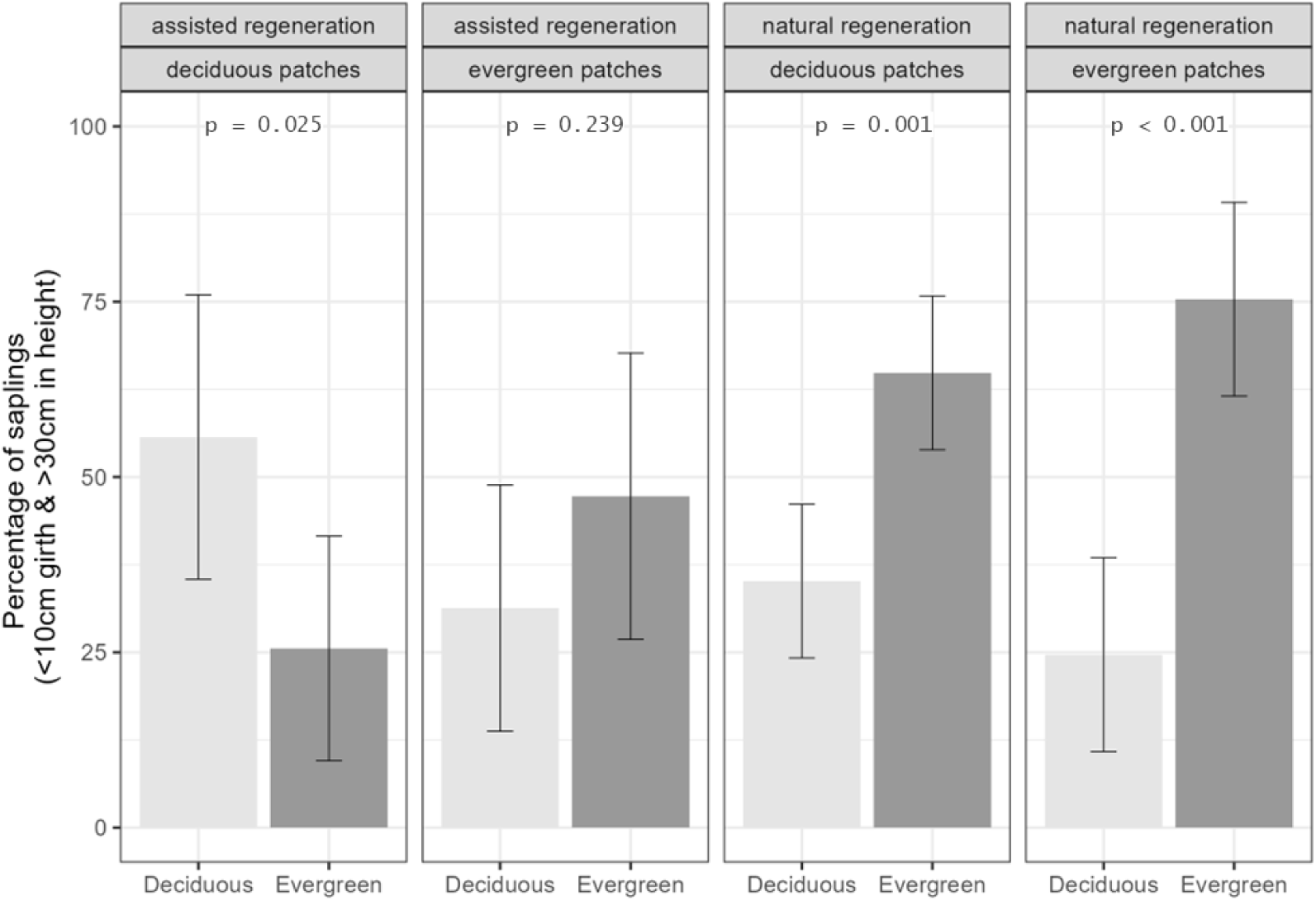
A comparison of the % of saplings that are either deciduous or evergreen disaggregated by regeneration type – natural regeneration or assisted regeneration through sapling planting – and by forest type, with thirty 0.01ha plots in deciduous (n=16) or evergreen (n=14) patches. P-values from statistical tests of differences in fraction of evergreen versus deciduous habit saplings are from a parametric t-test.

Our results on large tree density and diversity indicate that logging intensity has likely decreased after 2005, in line with the working plan. We make this inference because forests logged under the old policy in the 1990s had thrice as much time to recover as forests logged in the 2007-14 period, and yet when we enumerated standing timber stock in 2018, we found little or no differences in overall density and species richness. The new policy is in line with extraction quotas typical of reduced-impact logging practices, and these practices are well understood to facilitate forest recovery (Roopsind et al. 2018). Simultaneously, the seemingly poor recovery of older logged forests (where the intensity of logging was very high) corroborates official records of very high timber extraction under the old policy (Sekhsaria 2004) - an observation that forest staff across the archipelago made throughout the project. However, these broad summaries hide species-level changes of relevance to timber extraction.

Rank abundance curves reveal a long-term negative effect of logging operations on several dominant species. A 33% and 50% decline in large Padauk trees after one and two logging events respectively (Figure 3) highlight both the preferential harvest of Padauk and its relatively slow growth rates (Prasad and Dutt 2008). Interestingly, species like Padauk (*Pterocarpus dalbergioides*), Papita (*Pterocymbium tinctorium*) and Tompeing (*Artocarpus chaplasha*) with a predilection for semi-evergreen or deciduous patches do occur within evergreen patches, as seen in the rank-abundance curve for unlogged evergreen patches (Figure 3). Here, because understory light is more limiting, these trees of deciduous habitat develop taller, straighter boles than their counterparts in more open deciduous patches (Loubota Panzou et al. 2021). Naturally, these high-value species are preferentially removed from evergreen patches. This, in combination with their reduced rate of regeneration in evergreen habitats (Figure 4) contributes to their rarity in evergreen habitats (Figure 3).

In contrast, Garjan (*Dipterocarpus sp.*) trees, found exclusively within evergreen forests (except *D. incanus*, but *D. incanus* are rare in this study region) appear to be resilient to low-intensity logging, their abundance even increasing as forests that are logged once and then twice (Figure 3). While these do point to high growth rates and a stable growing stock of Garjan (*Dipterocarpus sp.*) -- a well-recognized strategy of advanced regeneration among Dipterocarps across South-East Asia (Ghazoul 2016) -- the same trend may also be the result of the local extinction of Andaman wild pigs (*Sus scrofa*) within worked forests for the last several decades (pers. comm., field staff). Across southeast Asia, pigs play a crucial role in controlling the dominance of dipterocarps by tracking fruiting individuals and eating their seeds (Ickes 2001). While fungi and insects can compensate for the pigs’ role to some extent (Williams et al. 2021), high light and low moisture in the understory of logged forests limit their effectiveness (Pillay et al. 2018). Throughout the study period of 8 months, we only found three signs of digging activity of wild pigs, and no signs were found in twice-logged forests.

Analysis of the first dataset reveals that the new working plan has achieved partial success in allowing forest recovery by lower extraction quota. However, stark differences in recovery between evergreen and deciduous patches, as well as differences in dominant species within each of these patches in response to logging point to an urgent need to examine forest recovery in these two forest types separately (Poorter et al. 2019). Our data on regeneration reveals that significant changes are required in assisted regeneration activities if forest management is to align with a policy of eco-restoration, forest reclamation, biodiversity conservation and support of local livelihoods.

### Recommendations

Putting together the results from our study and literature as a whole, we make the following recommendations to improve forest management under the current working plan:

#### 1. Delineation of coupes based on forest type, and not convenience alone

It is easy to clearly identify evergreen and moist-deciduous forests during the leaf-off summer months (January to March) using freely-available satellite imagery such as NDVI/EVI images from LANDSAT data. Coupe maps can be drawn over and above this satellite composite, separately setting out coupes for each of the two forest types. We recommend that semi-evergreen forests be conservatively treated as evergreen forests because, in our limited experience, it is difficult to separate highly degraded evergreen forests from intact semi-evergreen forests.

#### 2. Reduced extraction quotas for deciduous forests

Deciduous forests world-over recover slowly (Poorter et al. 2019) and Andaman Islands is no exception. Either through separate coupes (see #1 above) or otherwise, removing fewer deciduous habit trees within deciduous patches (for all canopy species, but especially *padauk* (Prasad and Dutt 2008), to recover) and evergreen patches (thereby reducing *garjan* dominance) will greatly benefit biodiversity, carbon and long-term sustainability of timber stocks in the region.

#### 3. Reorder sequence of logging of coupes within a felling series to maximize recovery

The current working plan is a hard reset in 2005 and thus, logging coupes include variously logged forests: those that have barely recovered (like the forests of Nilambur, the twice-logged treatment with less than 20 years in between) to forests with a very long recovery period (like the Bajalungta forests that were previously logged in the 1940s, the recent once-logged treatment). This information can be used to reorder the sequence of logging of coupes within a given felling series to maximize recovery: coupes that were logged the latest can be logged at the end, while coupes logged long ago can be the first to be worked.

#### 4. Minimizing forest disturbance from tending and culture operations with selective logging to mitigate rising deciduousness

Forest disturbance across the world increases deciduous elements in the forest (Biswas et al. 2024) and the Andaman Islands is no exception. Logging-related increase in deciduous elements was first noted by botanists Shanti Nair and Satish Nair in 1983: they speculated that tending and culture operations associated with the Andaman Shelterwood System such as burning, weeding and cleaning were eliminating evergreen species from deciduous patches (Sekhsaria and Pandya 2010). A larger fraction of deciduous habit species is also commercially viable– a phenomenon confirmed by a study from South Andaman (Saha 1992). In 2003-05, the Shekhar Singh commission report cited these two sources to recommend an overhaul in timber extraction to an approach that is more mindful of biodiversity and forest recovery. Our previous study (Surendra et al. 2021b) confirmed that with increasing disturbance (twice logged versus once logged), the fraction of deciduous stems in the understory increased along tree-fall gaps created by logging (Figure 3c, Surendra et al. 2021). Together, we recommend that any tending practices that increase forest disturbance must be strictly avoided.

#### 5. Identify pockets of unlogged forests within logged territorial forests and protect them from future logging

We found unlogged patches within territorial forests that have been designated for logging, whereas our data corroborates their superior density and diversity of trees (Figure 2, 3). Such pockets of unlogged forests within logging concessions are invaluable for effective forest recovery (Gustafsson et al. 2012; Putz et al. 2019) and therefore must remain protected. We located these unlogged pockets using local knowledge of forest department field staff, especially older employees who remember logging history (unlogged forests are called “*khara jungal*” in the local Ranchi language): such local knowledge of unlogged patches must be leveraged at the divisional level to avoid future disturbance of these areas. Moreover, unlogged patches also included (a) areas that were within the Jarawa tribal reserve until recently (such as two evergreen unlogged plots on Evergreen “Island” in Baratang Forest Division), (b) infertile sites with poor soils where trees develop large buttresses and short stature with low timber value (such as plots in the northern-western fringe of Baratang island) and (c) pockets of forests with valuable timber but difficult to extract, like valley bottoms (two evergreen unlogged plots on Baratang Island of the rare Andaman Giant Evergreen Forest type) and steep hill slopes outside the ridge line. All these unlogged patches must be identified at local-scales and protected.

#### 6. An overhaul of assisted regeneration to plant saplings of species that improve biodiversity and livelihood outcomes, and not just timber

a. Deciduous saplings must be planted in deciduous patches alone, and not in evergreen sites (Figure 4) as planting deciduous species in evergreen patches will increase and not reverse deciduousness.
b. A majority of the saplings planted (pers. comm, forest staff) and all plots in this study were on a logging coupe road. When these coupes are logged again 30 years later as per the working plan, logging roads will be laid again, and these planted trees will need to be harvested. Thus, focussing tree planting in tree-fall gaps around logging stumps rather than coupe roads (with high natural regeneration anyway) might be more efficient.
c. Taller saplings survive more (Banin et al. 2022) and yet, the alignment of tree-planting regimes to annual cycles mean that the oldest saplings planted in the forest are at most 1 year old. Planting older (and therefore taller saplings) saplings on rotation will improve survival success
d. The selection of species for planting continues to be dominant species of timber value. In order to meet the mandate for biodiversity conservation and livelihood protection of the new working plan, we recommend a much wider set of species, excluding common timber species with ample natural regeneration (e.g., *Dipterocarpus gracilis, D. grandiflorus*) and including important species beyond timber. These include –

i. uncommon large-seeded canopy species such as *Garuga pinnata, Endocomia macrocarpa var. prainii* and *Aglaia spectabilis* with high ecological and carbon sequestration value
ii. ecologically unique species like *Corypha umbraculifera* (sensitive due to a semelparous life-cycle), *Podocarpus neriifolius, Mesua manii*, *Magnolia andamanica* (uncommon and slow-growing species) and *Pisonia excelsa* (trunk damaged by elephants, against which it has evolved no protection)
iii. trees of NTFP value such trees for fruits (e.g., *Baccaurea sapida*, *Spondias pinnata*), leaves (e.g., *Champereia manillana*) and small timber (several Annonaceae including *Sagraea elliptica* and *Pseuduvaria prainii*; coppicing tree *Hunteria zeylanica* that is used as a fence post).

To achieve this, we recommend systematic identification of mother trees (preferably in disturbed forests), collection of seeds and development and codification of nursery techniques.

Table 1 provides a starting point with a list of such species compiled from local information gathered throughout the study period. Moreover, forest departments and NGOs like the Nature Conservation Foundation in India and across tropical forest countries around the world have developed guidelines and best practices to restore rainforest tree species irrespective of their timber value (Gunatilleke et al. 2023; Osuri et al. 2024). We recommend training sessions by such practitioners to augment substantial local knowledge within the forest department staff.

**Table 1:**
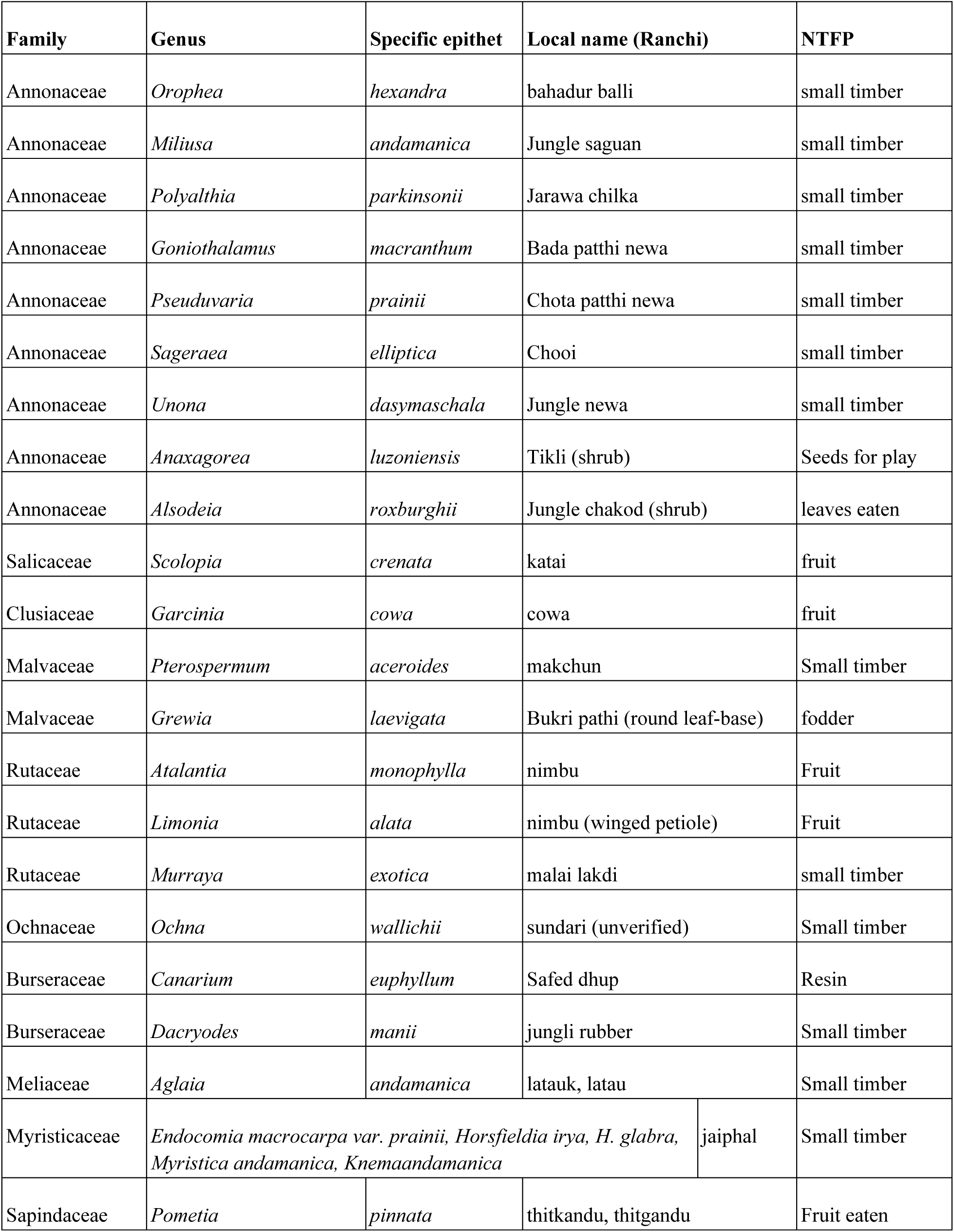
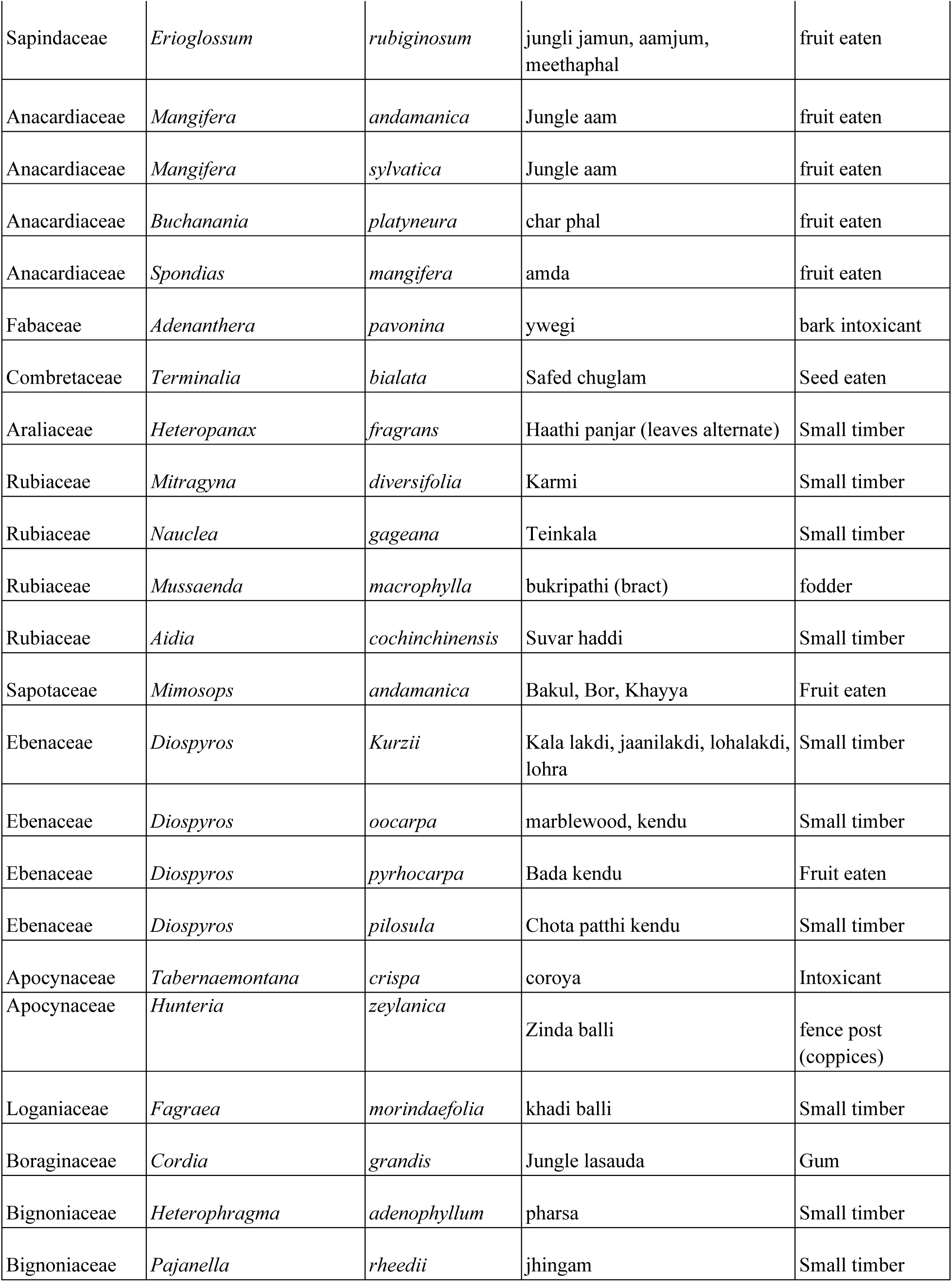
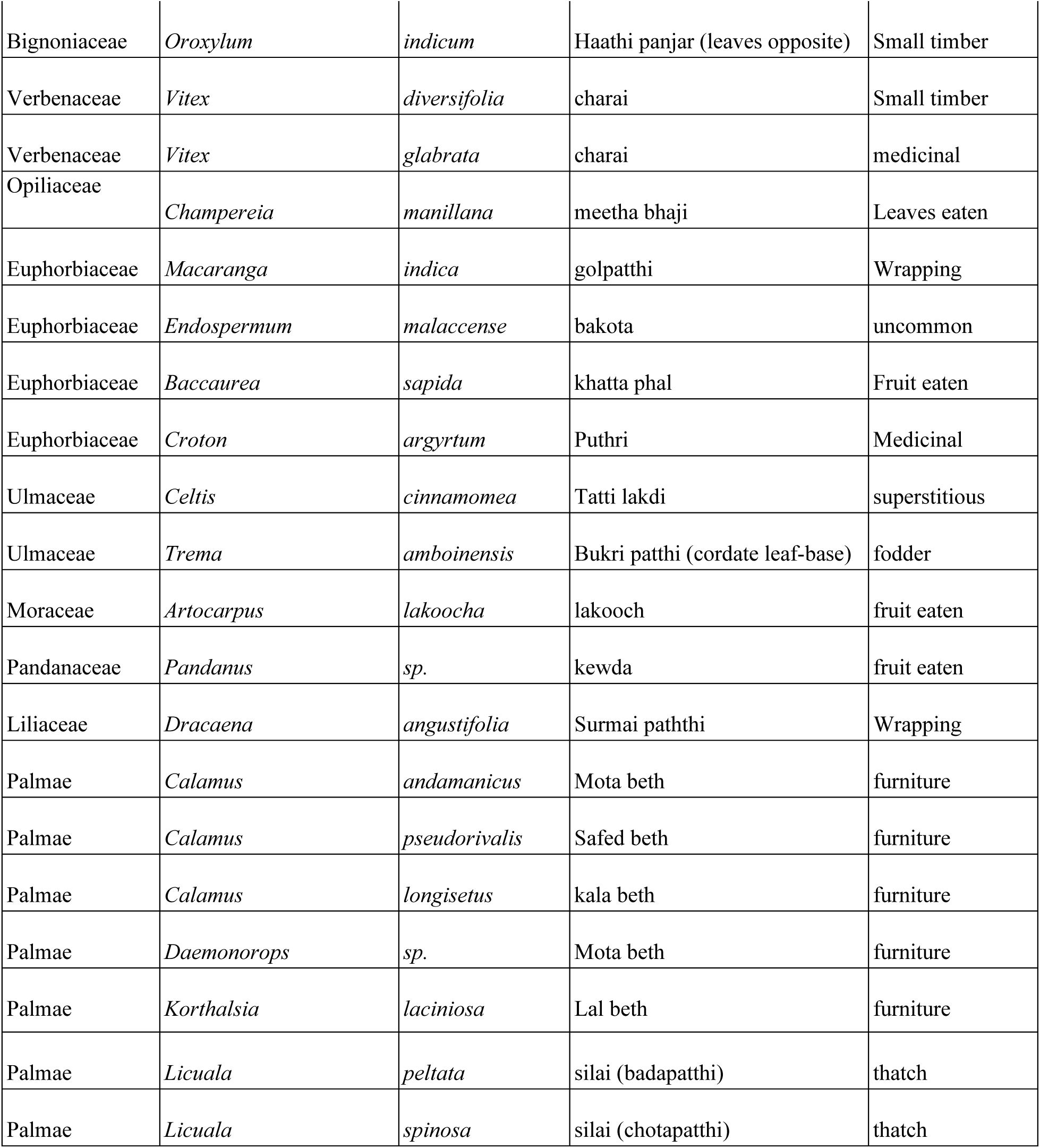
Non-exhaustive list of trees and palms used as non-timber forest products: potential for inclusion in assisted regeneration under the Working Plan’s eco-restoration mandate.

## CONCLUSION

Our survey shows that logging intensity has reduced under the new policy, compared to the old policy, in line with the working plan and has resulted in retaining large trees in the forests under the new regimes. However, our study revealed strongly detrimental legacy effects of past logging regimes on the relative abundance of large timber trees. We identify three strategies -- identification and exclusion of unlogged forest patches within logging coupes, reordering of coupes’ logging sequence to maximize forest recovery, and separate extraction quotas for deciduous and evergreen forest types, in proportion to their acreage within each coupe -- to substantially improve forest recovery, in line with the working plan. Further, our data and field observations highlight inadequacies in assisted regeneration within the eco-restoration working circle and we make recommendations to address these gaps. We believe that this study, in combination with a previously published work with a separate but complementary dataset (Surendra et al. 2021b), provides a useful framework to make similar assessments of forest management in the future and improve selective logging and eco-restoration practices in line with the Working Plan (Singh 2003).

## ACKNOWLEDGEMENTS

We thank PCCF D.M. Shukla IFS, PCCF Tarun Coomar IFS and CCF (Territorial) M. Raj Kumar IFS for their continuous support, DFO Middle Andaman Jabestin Arulraj IFS for support with accommodation, logistics and for insightful discussions, and collaborator DFO Baratang Division Vanjulavalli Sridhar IFS for championing this project and facilitating a rare opportunity for critical and independent scientific assessment of forest management. We sincerely thank all ground-level staff in Baratang and Middle Andaman Island, members of the ANET team and residents of Baratang and Kadamtala who, through ways big and small, supported this project. We are especially grateful to field assistants Adhir Samadder, Jagadish Mandol, Ravi Mandol, Alexius Kujur, Binod ji, Niranthar ji, Edmond Kujur, Chandan, Lochan, Prafulla da and Kaushik for extensive support on and off field. We are grateful to Navendu Page and Joju Alappatt for help with tree identification, and Botanical Survey of India - Port Blair for providing access to herbarium samples to aid identification. This project was part of A. Surendra’s M.Sc. thesis and was funded by the Department of Atomic Energy, Government of India (TIFR) and the Tata Trust. AMO received fellowship support from the Indian Government’s Department of Biotechnology via a Ramalingaswami Re-entry Fellowship (Grant BT/RLF/Re-entry/58/2018).

## DECLARATION

I/We declare that the manuscript has not been published in any journal/book or proceedings or any other publication, or offered for publication elsewhere in substantially the same or abbreviated form, either in print or electronically.

## REFERENCES

Banin LF, Raine EH, Rowland LM, et al (2022) The road to recovery: a synthesis of outcomes from ecosystem restoration in tropical and sub-tropical Asian forests. Philosophical Transactions of the Royal Society B: Biological Sciences 378:1–17.

Bartholomew DC, Hayward R, Burslem DFRP, et al (2024) Bornean tropical forests recovering from logging at risk of regeneration failure. Global Change Biology 30:1–19.

Bhattee SS (1958) Logging in the Andamans. Indian Forester 84:197–212

Biswas N, Sadekar V, Biniwale S, et al (2024) Chronic disturbance of moist tropical forests favours deciduous over evergreen tree communities across a climate gradient in the Western Ghats. bioRxiv, 2024–01

Blanc L, Echard M, Herault B, et al (2009) Dynamics of Aboveground Carbon Stocks in a Selectively Logged Tropical Forest. Ecological Applications 19:1397–1404

Chaudhary A, Burivalova Z, Koh LP, Hellweg S (2016) Impact of Forest Management on Species Richness: Global Meta-Analysis and Economic Trade-Offs. Scientific Reports 6:1–10

Chaudhry P (1998) Striking Features of Andaman Forestry. Indian Forester 124:463–472

Chengappa BS (1934) Andaman forests and their reproduction. Indian Forester 60:

Fisher B, Edwards DP, Giam X, Wilcove DS (2011) The high costs of conserving Southeast Asia’s lowland rainforests. Frontiers in Ecology and the Environment 9:329–334.

Ghazoul J (2016) Dipterocarp Biology, Ecology, and Conservation. Oxford University Press Gibson L, Lee TM, Koh LP, et al (2011) Primary forests are irreplaceable for sustaining tropical biodiversity. Nature 478:378–381.

Gunatilleke N, Neidel JD, Raman TRS, et al (2023) Ecological Approaches to Forest Restoration: Lessons Learned from Tropical Wet Asia. In: Florentine S, Gibson-Roy P, Dixon KW, Broadhurst L (eds) Ecological Restoration: Moving Forward Using Lessons Learned. Springer International Publishing, Cham, pp.103–147

Gustafsson L, Baker SC, Bauhus J, et al (2012) Retention Forestry to Maintain Multifunctional Forests: A World Perspective. BioScience 62:633–645.

Ickes K (2001) Hyper-abundance of Native Wild Pigs (Sus scrofa) in a Lowland Dipterocarp Rain Forest of Peninsular Malaysia1. Biotropica 33:682–690.

Krishnan A, Osuri AM (2023) Beyond the passive–active dichotomy: aligning research with the intervention continuum framework of ecological restoration. Restoration Ecology 31:1–6.

Loubota Panzou GJ, Fayolle A, Jucker T, et al (2021) Pantropical variability in tree crown allometry. Global Ecology and Biogeography 30:459–475.

Malhi Y, Gardner TA, Goldsmith GR, et al (2014) Tropical Forests in the Anthropocene. Annual Reviews in Environmental Resources 39:125–159.

Osuri AM, Kasinathan S, Raman TRS, Mudappa D (2024) Restoration opportunities beyond highly degraded tropical forests: Insights from India’s Western Ghats. Biological Conservation 291:1–6.

Pal A, Dutta S, Mukherjee PK, Paul AK (2005) Occurrence of heavy metal-resistance in microflora from serpentine soil of Andaman. Journal of Basic Microbiology 45:207–218.

Pal A, Paul AK (2007) Rhizosphere of Nickel hyper-accumulating plants: a niche for nickel-resistant bacteria. In: Roy A.K. (ed) Rhizosphere Biotechnology: Plant Growth Retrospect and Prospect, Scientific Publishing India, pp. 121–134

Parkinson CE (1923) Forest Flora of the Andaman Islands.

Pillay R, Hua F, Loiselle BA, et al (2018) Multiple stages of tree seedling recruitment are altered in tropical forests degraded by selective logging. Ecology and Evolution 8:8231–8242.

Poorter L, Rozendaal DMA, Bongers F, et al (2019) Wet and dry tropical forests show opposite successional pathways in wood density but converge over time. Nature Ecology & Evolution 3:928–934.

Prasad PRC, Dutt CBS (2008) Population Structure, Age Gradations, and Regeneration Status of Pterocarpus dalbergioides Roxb., An Endemic Species of Andaman Islands, India. The Pacific Journal of Science and Technology 9:658–664.

Putz FE, Baker T, Griscom BW, et al (2019) Intact Forest in Selective Logging Landscapes in the Tropics. Frontiers For Global Change 2:1–10

Putz FE, Sist P, Fredericksen T, Dykstra D (2008) Reduced-impact logging: Challenges and opportunities. Forest Ecology and Management 256:1427–1433

Putz FE, Zuidema PA, Synnott T, et al (2012) Sustaining conservation values in selectively logged tropical forests: the attained and the attainable. Conservation Letters 5:296–303.

R Core Team R (2018) R: A language and environment for statistical computing. 2018

Reddy CS, Satish KV, Pasha SV, et al (2016) Assessment and monitoring of deforestation and land-use changes (1976–2014) in Andaman and Nicobar Islands, India using remote sensing and GIS. Current Science 111:1492–1499

Roopsind A, Caughlin TT, van der Hout P, et al (2018) Trade-offs between carbon stocks and timber recovery in tropical forests are mediated by logging intensity. Global Change Biology 24:2862–2874

Saha S (1992) M.Sc. Thesis: Regeneration of important rainforest tree species in virgin and selectively logged sites in the South Andaman Island. Salim Ali School of Ecology, Pondicherry University

Sekhsaria P (2004) Illegal Logging and Deforestation in Andaman and Nicobar Islands, India. Journal of Sustainable Forestry 19:319–335

Sekhsaria P, Pandya V (2010) Jarawa Tribal Reserve dossier: cultural & biological diversities in the Andaman Islands. UNESCO

Singh MP (2003) Report: Working plan for South Andaman Forest Division (for the period from 2003–13). In compliance with the order of the Honorable Supreme Court dated 07.05.02. Andaman & Nicobar Administration, Department of Environment and Forests, Van Sadan, Port Blair, Andaman & Nicobar Islands

Surendra A (2023) Github Repository. https://github.com/akshaysurendra/andaman-logging-management

Surendra A, Osuri A, Ratnam J (2021a) Dataset: Varying impacts of logging frequency on tree communities and carbon storage across evergreen and deciduous tropical forests in the Andaman Islands, India. https://datadryad.org/stash/dataset/doi:10.5061/dryad.1c59zw3rp

Surendra A, Osuri AM, Ratnam J (2021b) Varying impacts of logging frequency on tree communities and carbon storage across evergreen and deciduous tropical forests in the Andaman Islands, India. Forest Ecology and Management 481:1–11

Williams PJ, Ong RC, Brodie JF, Luskin MS (2021) Fungi and insects compensate for lost vertebrate seed predation in an experimentally defaunated tropical forest. Nature Communications 12:1–8

Zimmerman BL, Kormos CF (2012) Prospects for Sustainable Logging in Tropical Forests. BioScience 62:479–487

